# FACS-assisted single-cell lipidome analysis of phosphatidylcholines and sphingomyelins

**DOI:** 10.1101/2022.02.15.479799

**Authors:** Sarah E Hancock, Eileen Ding, Emma Johansson Beves, Todd Mitchell, Nigel Turner

## Abstract

Recent advances in single-cell genomics and transcriptomics technologies have transformed our understanding of cellular heterogeneity in growth, development, ageing and disease; however, methods for single-cell lipidomics have comparatively lagged behind in development. We have developed a high-throughput method for the detection and quantification of a wide range of phosphatidylcholine (PC) and sphingomyelin (SM) species from single cells that combines fluorescence-assisted cell sorting (FACS) with automated chip-based nanoelectrospray ionization (nanoESI) and shotgun lipidomics. We show herein that our method is capable of quantifying more than 50 different PC and SM species from single cells and can easily distinguish between cells of different lineages or cells treated with exogenous fatty acids. Moreover, our method can detect more subtle differences in the lipidome between cell lines of the same cancer type. Our approach can be run in parallel with other single-cell technologies to deliver near-complete multi-omics data on cells with a similar phenotype and has the capacity to significantly advance our current knowledge on cellular heterogeneity.

## Introduction

Single-cell analysis has generated much interest in recent years and has revolutionized our understanding of the fundamental processes underlying cell biology, stemness, and clonality. Alongside this, single-cell research has provided astonishing new insight into the role of cellular heterogeneity in cancer, ageing, and other diseases. This work has largely been driven by the development of methods for transcriptomic profiling of single cells that use amplification strategies to augment analyte signal. In contrast, methods for assaying small molecules such as lipids from single cells have lagged in development due to the high complexity of lipids within cells (i.e., 1000s of different molecular species), their low abundance (*e*.*g*., amol to low fmol range), and an inability to increase their signal via amplification. However, we know that changes in lipid composition are significant in fundamental biological processes such as growth, development, differentiation, and ageing and are also important in disease states such as cancer. Being able to detect and quantify lipid composition in single cells would provide enhanced data on such cell states and reveal new information on the cellular role(s) of lipid heterogeneity.

Mass spectrometry (MS) is the current technology of choice for the comprehensive profiling of lipids from biological tissues due to its broad specificity and sensitivity in detecting very low amounts of analyte from biological samples (*e*.*g*., fmol to pmol range). MS is also emerging as a key technology in the detection of lipids from single cells; however, there are some limitations in the sensitivity and/or specificity of currently available technologies. Commonly used ionization techniques for lipid single cell analysis include matrix-assisted laser desorption ionization (1–10), secondary ion mass spectrometry (11–13), laser ablation electrospray ionization (14, 15), nanospray desorption electrospray ionization (16) and nanoelectrospray ionization (NanoESI) (17–28). NanoESI capillaries are an ideal size for directly sampling and infusing the contents of a single cell into the mass spectrometer, but this process is quite laborious and requires the use of micromanipulators for handling and sampling from single cells which limits its throughput. Some improvements to the throughput of the technique have been made through the use of a continuous sampling arm that uses dual-bore quartz needles to sample from cells located on a movable stage (*i*.*e*., single probe MS) (21–23, 26–28). However, the spray duration per cell is limited (∼3 min) which precludes the acquisition of tandem mass spectrometry data from all but the most abundant metabolites (26). Improvements in spray duration from a single cell have been made by reducing the flow of analyte into the instrument (*i*.*e*., picoESI) (24). Using this method Wang and colleagues were able to characterize several biomarkers for different breast cancer cell subtypes, including membrane lipid species (24).

In this work we sought to develop an MS-based single-cell lipidomics workflow that is both high throughput and quantitative. To that end, we combined single cell isolation by fluorescence-assisted cell sorting (FACS) with shotgun lipidomics and a chip-based nanoESI ionization source. This nanoESI source is fully automated and has been used previously for single cell assay using its liquid-extraction surface analysis (LESA) operating mode (17, 18). In contrast, we use direct infusion mode to avoid ambient lipid oxidation following the printing of cells onto a surface for analysis for LESA (29) and sorted cells into a 96 well plate to increase throughput. We then used a semi-targeted shotgun lipidomics approach that could facilitate both improved sensitivity and provide broad specificity in detecting a range of lipid species that may be present both similar and different cell populations. Using this approach we have been able to detect and perform relative quantitation on more than 50 discrete phosphatidylcholine (PC) and sphingomyelin (SM) species from single cells isolated from different human cell types, and we demonstrate the utility of our method for detecting potentially clinically relevant heterogeneity in prostate cancer cells.

## Materials and Methods

### Cell culture

HepG2 and C2C12 cells were obtained from American Type Cell Culture Collection (ATCC; Manassas, Virginia, USA), while prostate cell lines (LNCaP, PC3 DU145, and PNT1) were gifted by Dr Andrew Hoy (University of Sydney Australia). HepG2 and C2C12 cells were cultured in high-glucose DMEM supplemented with 10% heat-inactivated fetal calf serum (FCS; Sigma-Aldrich Pty Ltd, Sydney, NSW, Australia) incubated at 37ºC in 5% CO_2_. Prostate cancer cell lines were cultured in RPMI medium supplemented with 2 mM L-glutamine and 10% heat-inactivated FCS (ThermoFisher Scientific, Melbourne, VIC, Australia). Cell media was changed every three days and cells were passaged regularly at ∼80-90% confluency by trypsinization. Cell lines were regularly screened for mycoplasma infection. Cell number was determined by trypan blue staining using an automated cell counter (Invitrogen™ Countess II, ThermoFisher Scientific, Melbourne, VIC, Australia).

### DHA supplementation

DHA was obtained from Avanti Polar lipids (Alabaster, AL, USA) and dissolved in 100% ethanol at a concentration of 100 mM. DHA was conjugated to fatty acid-free BSA (2% w/v; Sigma-Aldrich Pty Ltd, Sydney, NSW, Australia) in media to increase both its solubility and bioavailability (30). 25 μL of DHA stock or 25 μl of 100% ethanol (CON) was added per 50 mL of normal growth media containing 2% (w/v) of fatty acid-free BSA. Media was sterile filtered by passing it through a 0.2 μM PES filter, and then both CON and DHA-containing media were incubated in a water bath for 2 hours at 55ºC to conjugate DHA to BSA. Conjugated media was stored at 4ºC until used. Cells were incubated overnight in the presence of pre-warmed CON or DHA-conjugated media before harvest.

### Bulk cell lipid extraction

All solvents and additives used for lipid extraction were of the highest grade available (either LCMS or HPLC grade; ThermoFisher Scientific, Melbourne, VIC, Australia). Bulk cell extracts were prepared from adherent cells grown in 6 well plates. C2C12 cells were seeded at 1×10^5^ cells/well and HepG2 at 2×10^5^ cells/well and grown to ∼70-80% confluency (∼2-3 days). Cells were then treated with CON or DHA-containing media and harvested the following day (final confluency ∼80-90%). Media was removed from wells and cells were washed with ice-cold phosphate-buffered saline (PBS, pH 7.4). Methanol containing 0.01% butylated hydroxytoluene (BHT; 300 μl) was added to each well, and plates were scraped into 2.0 mL eppendorfs. An additional 100 μl of methanol was used to wash each well/scraper and was combined with cell extract. Empty wells were also scraped for background normalization. To the methanolic cell extracts, 920 μl of methyl tert-butyl ether was added, and samples were vortexed at 1200 rpm for 2-3 hours at room temperature (MixMate ®, Eppendorf South Pacific, Sydney, NSW Australia). Following this, 230 μl of 150 mM ammonium acetate was added and samples were vortexed vigorously for a minimum of 30 seconds. Samples were centrifuged for 5 mins at 2,000 *x* g to ensure phase separation and the upper organic phase was removed to a 2 mL glass vial. This organic phase was then diluted 500-fold in methanol:chloroform (2:1 v/v) containing 5 mM ammonium acetate and stored at -30ºC until analysis. Prior to analysis 40 μl of diluted cell lipid extract was added to a 96 well plate, which was then sealed.

### FACS and lipid extraction

HepG2 and C2C12 cells were grown in 75 cm^2^ flasks until ∼70-80% confluent before being treated overnight with CON or DHA media. The following day cells (∼80-90% confluency) were trypsinized and counted. Prostate cells (LNCaP, DU145, PC3 and PNT1) were grown to ∼80-90% confluency before harvest by trypsinization followed by cell counting. 1-2 million cells of each line were centrifuged (300 *x* g, 5 min) and resuspended in PBS containing 10% FCS and 2 mM EDTA. This process was then repeated to remove all traces of growth media. Cells were placed on ice and sorted within an hour by FACS (BD FACSAria™ III, BD Biosciences, Sydney, NSW Australia) directly into a 96-well plate preloaded with methanol spiked with 0.01% BHT and internal standards (PC 17:0/17:0 and dihydrosphingomyelin, DHSM, 12:0; Avanti Polar Lipids, Alabaster, AL, USA). Internal standards were added at a rate of 1000 fmol per well for fifty cells and 1.38 fmol per single cell. Plates were sealed and stored at -80ºC until analysis. Prior to analysis methanol:chloroform containing 5 mM ammonium acetate was added to each well to achieve a final ratio of 2:1 v/v (final volume 40 μl). Wells containing solvent and internal standard only were included and used for background subtraction (*i*.*e*., extraction blanks).

### Mass spectrometry and lipid identification

NanoESI mass spectrometry of lipid extracts was performed using a hybrid triple quadrupole linear ion trap mass spectrometer (QTRAP® 5500, SCIEX, Framingham, MA, USA) equipped with an automated chip-based nanoelectrospray source (TriVersa Nanomate®, Advion Biosciences, NY, USA). Spray parameters were set at a gas pressure of 0.4 psi and a voltage of 1.2 kV. PC and SM data were acquired in positive ion mode using a precursor ion scan of *m/z* 184 at a scan rate of 200 Da/s across a mass range of 640 – 850 *m/z*. Declustering potential was set at 100 V, entrance potential at 10 V, collision energy at 47 V, and collision cell exit potential at 8V (31). Aspiration of 10 μl of sample from each well generated a stable spray time of ≥30 minutes.

Lipids were identified from acquired data using Lipidview™ software (v1.2b, SCIEX, Framingham, MA, USA). Processing settings in Lipidview™ were set at a mass tolerance of 0.5 Da, with a minimum intensity of 0.1% and a minimum signal-to-noise ratio of 4. Smoothing and deisotoping of lipid species were enabled. Lipid species were identified from target lists (see Supporting Information Table S1), and peak area for each detected lipid species was then exported. These data underwent further processing in R (32), including background subtraction and relative quantification from internal standards. Lipid nomenclature follows recommendations for the level of molecular detail known (33); and in the present study we report lipids as class (*e*.*g*., PC or SM) followed by the total number of carbons and carbon-carbon double bonds present within the fatty acids separated by a colon (*e*.*g*., a PC with 34 carbons and 1 double bond as PC 34:1). At the level of identification available by the technique used in this study some ambiguity exists between isobaric PC species containing either odd-chain or ether-linked fatty acid species, and in the absence of further structural detail we chose to report such species as ether-linked only where overlap exists.

### Statistical analysis

Statistical analysis was conducted in R (32), with t-distributed stochastic neighbor embedding (t-SNE) being performed using the Rtsne package (34). Parameters for t-SNE were set at default values except for perplexity which was set at ∼*N ^* (1/2) where *N* is the number of samples analyzed. Comparison between the relative intensity of PC and SM species detected from bulk cell extracts and fifty sorted cells was performed using an unpaired t-test with Welch’s correction, with statistical significance set at *P* < 0.05.

## Results

### Detection of phosphatidylcholine and sphingomyelin from single cells

To develop our single cell lipidomics method we used both immortalized myocytes (C2C12) and hepatocarcinoma cells (HepG2) cultured under normal growth conditions. To test our methods sensitivity we first used FACS to sort three lots of 50 cells from both cell lines and acquired baseline precursor *m/z* 184 scans to identify both PC and SM species. The resulting mass spectra for C2C12 cells are shown in Figure 1A. The data demonstrate a typical profile of both PC and SM species normally detected within C2C12 cells, with PC 34:1 (760.6 *m/z*) being the most abundant endogenous species present. Importantly, both PC and SM species are being detected at a similar intensity to that seen with cell extracts obtained using a standard extraction protocol before analysis by shotgun lipidomics (described below). We next infused up to 18 singly sorted C2C12 or HepG2 cells and acquired the same precursor *m/z* 184 scan, with representative spectra from 7 randomly selected single C2C12 cells shown in **Error! Reference source not found**.B. Not surprisingly there was a substantial increase in noise within our single-cell spectra; however, both the relative intensity and masses of each species detected closely match that obtained from the fifty cell samples (**Error! Reference source not found**.A). We calculated the signal-to-noise (S/N) for several PC and SM species at differing intensities (shown in **Error! Reference source not found**.B) demonstrating that the intensity of many of the detected PC and SM species are well above the background noise present.

**Figure 1:**
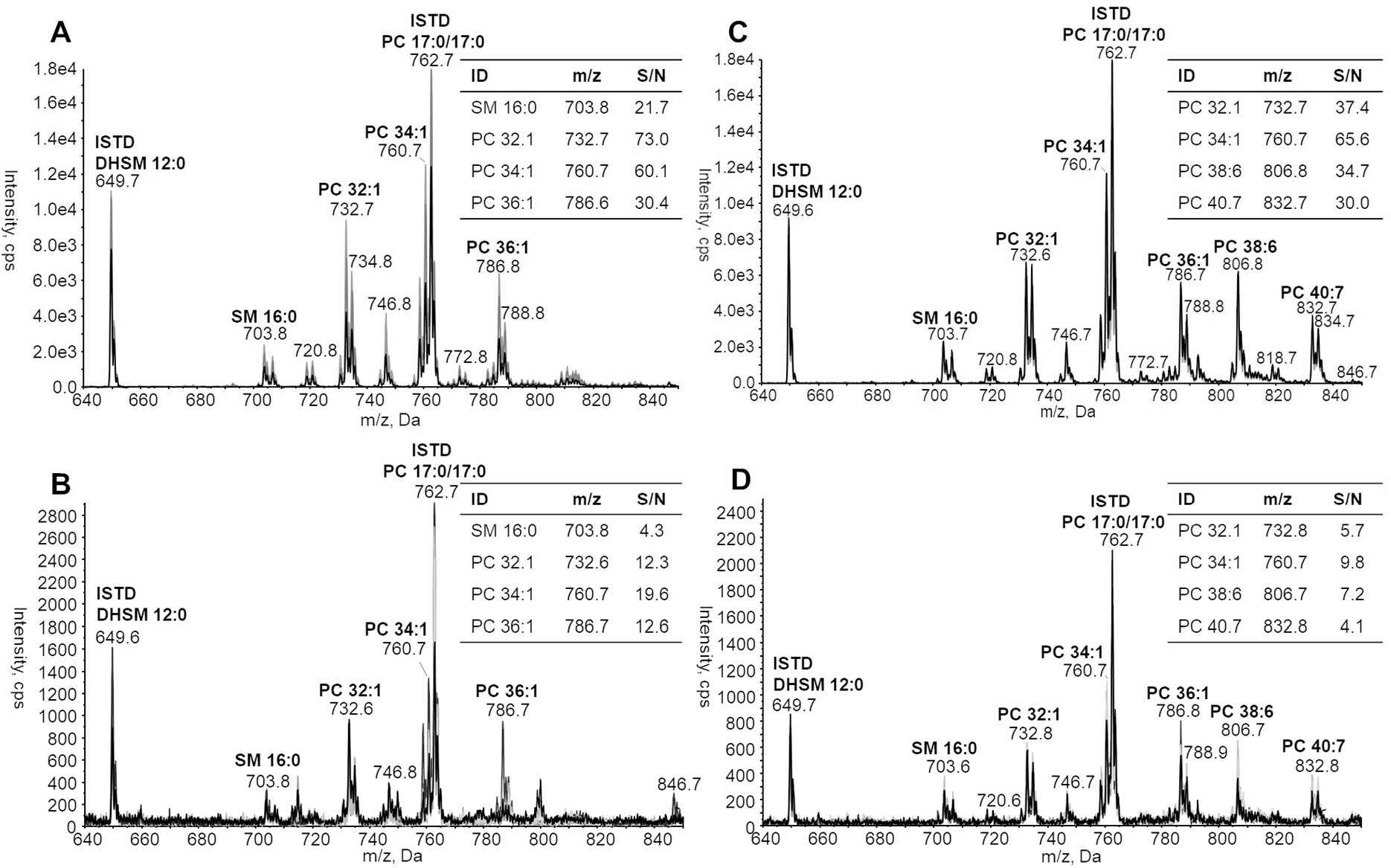
Precursor ion scan of *m/z* 184 obtained from fifty (A&C, n=3 biological replicates) and singly isolated (B&D, n=7) C2C12 cells obtained by fluorescence-assisted cell sorting. Cells were grown under normal culture conditions (A&B) or cultured overnight in the presence of 50 μM docosahexaenoic acid (DHA, C&D). Signal-to-noise (S/N) was calculated using the S-to-N Script in Analyst (v1.6.3, SCIEX, Framingham, MA, USA). Internal standards (ISTD) phosphatidylcholine (PC) 17:0/17:0 & dihydrosphingomyelin (DHSM) 12:0 were added at a concentration of 1000 and 1.38 fmol for fifty and single cells respectively.

To confirm that our method was accurately detecting PC and SM species from single cells we performed a second set of experiments in which our C2C12 cells were cultured overnight in the presence of 50 μM DHA. Given that FCS lacks meaningful levels of omega 3 (*i*.*e*., n-3) containing polyunsaturated fatty acids (PUFA) and human cells lack both the Δ12 and Δ15 desaturases necessary to create n-3 omega fatty acids and have limited capacity to elongate long-chain PUFA such as linolenic acid (18:3n-3) (35) we can be confident that any detected increase in DHA-containing PC species (*i*.*e*., species containing ≥6 carbon-carbon double bonds) can be directly attributed to the experimental conditions and thereby validate our single-cell lipidomics workflow. Overnight culture of C2C12 cells in the presence of DHA indeed produced the expected increase in DHA-containing PC species in both the fifty and single-cell spectra, including noticeable increases in PC 38:6 (806.8 *m/z*), PC 40:7 (832.7 *m/z*), and PC 40:6 (834 *m/z*) (**Error! Reference source not found**.C&D respectively). This increase in DHA-containing PC species within the single-cell spectra of DHA-treated cells confirms that we are measuring real biological information that is above the noise of the instrument, and together these data establish the validity of our approach for detecting biological variability in the lipids of single cells. All lipid species detected and quantified within single C2C12 and HepG2 cells are shown in Figure 2A&B respectively, including those detected under normal culture conditions (CON) and those from media supplemented with DHA. Overall we were able to detect 56 distinct PC and SM species from CON and DHA-treated single HepG2 and C2C12 cells. The inclusion of internal standards meant that we could perform relative quantitation on both PC and SM species detected from single cells, with the values reported being well within the expected range (16).

**Figure 2:**
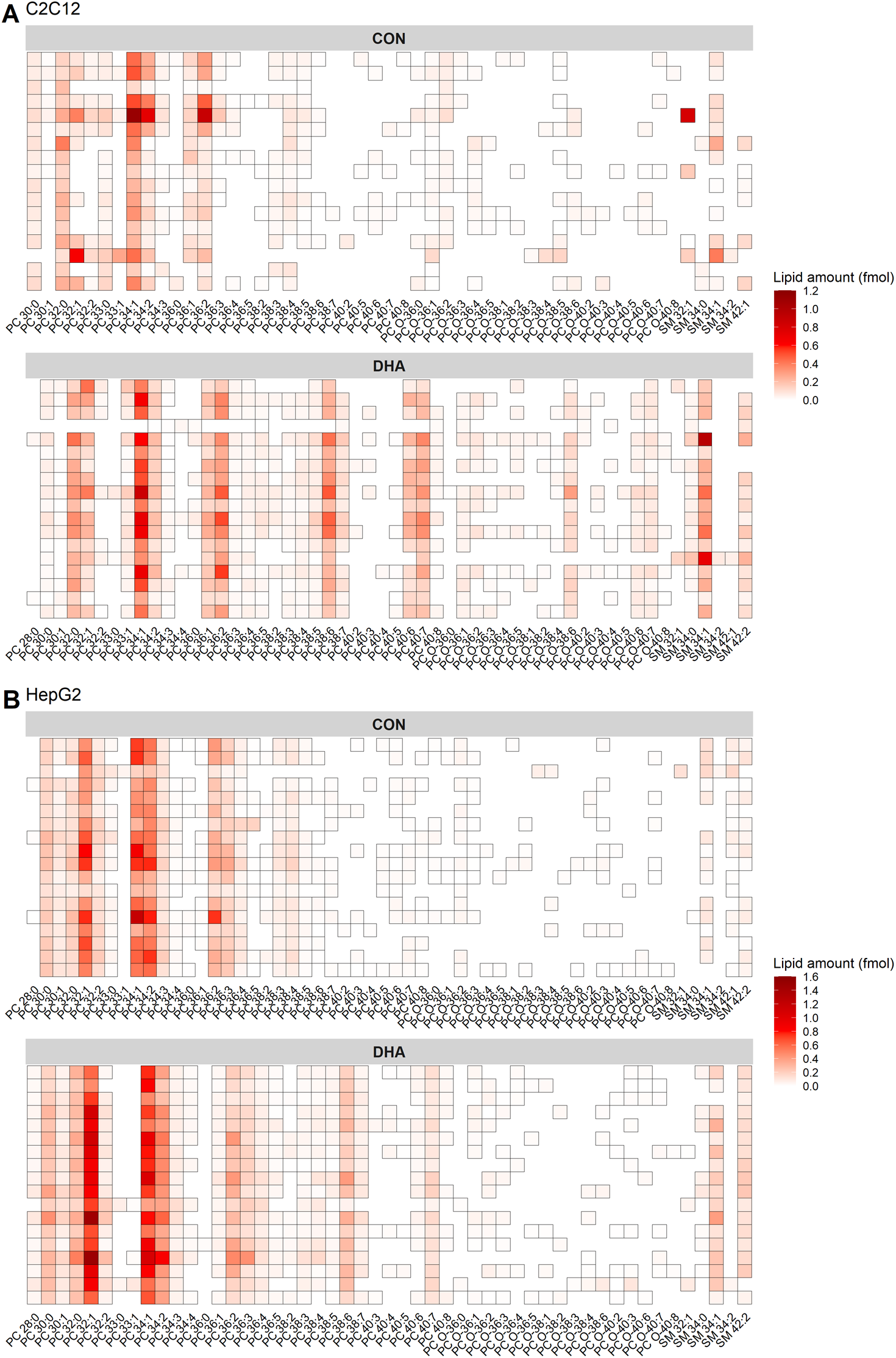
Heatmaps of phosphatidylcholine (PC) and sphingomyelin (SM) species detected in single isolated A C2C12 and B HepG2 cells. Cells were cultured under normal conditions (CON) or after overnight culture with 50 μM docosahexaenoic acid (DHA). Lipids were quantified from internal standards as described under method details and are present as relative quantified amount (fmol).

These data not only demonstrated a clear shift in lipid profile in DHA-supplemented cells, but we were also able to observe some inherent heterogeneity in PC and SM species within each of the cell lines and growth conditions. To further explore this heterogeneity we performed t-distributed stochastic neighbor embedding (t-SNE; **Error! Reference source not found**.) on our single-cell lipidomics dataset. This analysis showed clear clustering of data from each cell and treatment type, demonstrating that our method can discriminate easily between single cells from both different lineages and culture conditions. Also apparent within the data was some heterogeneity within a given cell line/ fatty acid treatment, which is demonstrated particularly within control HepG2 cells by the spread of the data across the second dimension of the t-SNE plot. Taken together, these data suggest that our single-cell lipidomics method can detect heterogeneity between cells derived from the same lineage but also can easily distinguish differences in the lipid profile of single cells isolated from different cell lineages and those enriched in exogenous lipid species.

### FACS has very little effect on cell PC & SM profile

As FACS is known to change the polar metabolite profile of cells (36) we next assessed its effects on PC and SM species composition. To that end we compared the relative intensities of bulk lipid extracts prepared using the traditional method of lipid extraction from adherent cells (*i*.*e*., bulk cell extract) to that of fifty cells isolated by FACS grown in either control or DHA-supplemented media (**Error! Reference source not found**.). No statistically significant differences in the relative intensity of PC or SM species were detected between bulk cell extracts or fifty-sorted C2C12 cells grown in control media, nor were any differences detected between bulk or fifty-sorted HepG2 cells grown in either control or DHA-containing media. Some small but statistically significant differences were observed between bulk extract and fifty-sorted DHA-supplemented cells for a few relatively minor PC & SM species (Supporting information, Table S2); however, these are extremely minor components of the cell membrane. Together these data suggest that FACS has very little impact on PC or SM profile, which further supports the validity of our methodological approach.

### Single-cell lipidomics can distinguish between different prostate cell lines

In the final part of this work we sought to validate our single-cell lipidomics workflow against an experimental condition with more subtle, yet potentially clinically relevant biological heterogeneity. We chose to use prostate cancer as our model for several reasons. First, it is widely reported that PC composition varies between different immortalized prostate cancer cell lines (37–39). Most recently Young and colleagues reported discrete differences in sum compositional PC species present across six different prostate cancer cell lines (38). For example, PC 34:1 was reported at ∼50% of total PC within the LNCaP prostate cancer cell line, whereas the PC3 cell line purportedly contained ∼30% PC 34:1 (38). Furthermore, applying single-cell lipidomics techniques to the study of prostate cancer is likely to yield clinically relevant information. Elevated expression of fatty acid synthase is common in metastatic, castration-resistant prostate cancer, and higher levels of phospholipids containing saturated and monounsaturated fatty acids are linked with greater metastatic potential in both prostate and other cancer types (reviewed by Butler et al.(40)). Acquired docetaxel resistance in prostate cancer cells can also increase overall PC and SM levels as well as the levels of some specific PC species (39). For these reasons prostate cancer was chosen as a model for our “proof-of-principle” experiments.

To that end we performed shotgun lipidomics on four different prostate cell lines sorted by FACS: three cancerous (DU145, LNCaP & PC3) and a non-tumorigenic immortalized prostate epithelial cell line (PNT1). All cell lines were grown in the same media and were approximately the same passage number. We initially confirmed the heterogeneity reported in the literature PC species between the different prostate cancer cell lines by analyzing fifty sorted cells (n=3 per cell line). The resulting PC profile proved to be very similar to that described by Young et al. (38), where PC 34:1 was present at ∼48% of total PC in LNCaP cells and ∼26% of PC within PC3 cells (**Error! Reference source not found**.A). We next performed single-cell lipidomics analysis on the four prostate cell lines (n=18 cells per line) and were able to detect just over 50 discrete PC and SM species in total from single cells (**Error! Reference source not found**.B). Not surprisingly, the overwhelming pattern of lipid species abundance in single cells closely mirrored that of the fifty cells data with some heterogeneity being apparent within discrete cells of each line. Finally, we performed t-SNE on our prostate cell single-cell lipidomics dataset, producing the plot shown in **Error! Reference source not found**.C. These data broadly cluster into three groups consisting of LNCaP, PC3, and a combination of DU145 and PNT1 cells. This suggests that there is some similarity in lipid profile between DU145 and PNT1 cells that is present even at the single-cell level. Increased heterogeneity appears to be present across the whole prostate single-cell dataset when compared with our previous model of two cell lines of different lineages treated with exogenous fatty acids as evidenced by the greater spread of data across both dimensions (*i*.*e*., Figure 3 vs **Error! Reference source not found**.C). Despite this increased variability, however, the data still show relatively tight clustering within the three broad groups of prostate cancer cell lines. This provides evidence that our single-cell lipidomics workflow is powerful enough to detect more subtle biological variation in cellular heterogeneity such as that found across different cancer cell lines.

**Figure 3:**
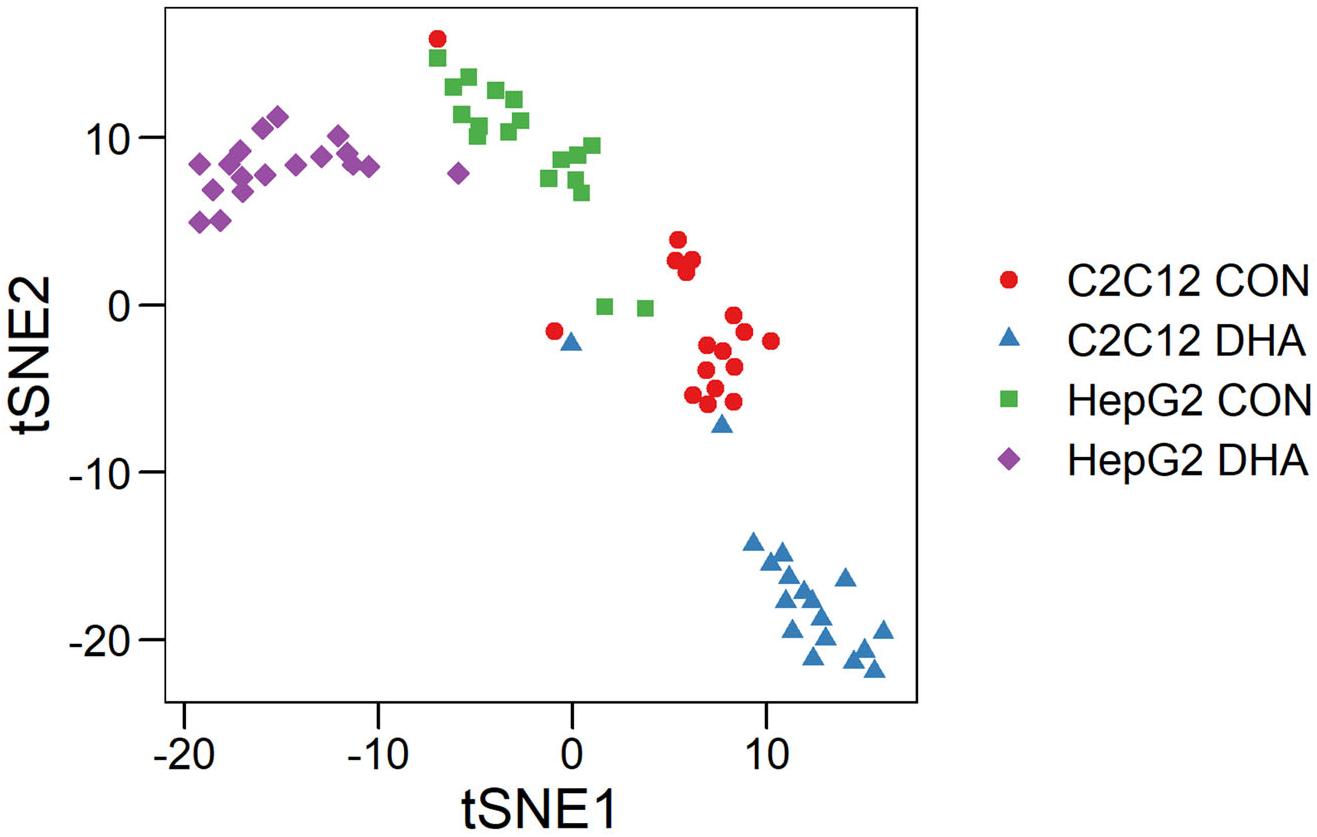
t-distributed stochastic neighbor embedding (t-SNE) of all single-cell data from both C2C12 & HepG2 cells grown in both control (CON) and docosahexaenoic acid (DHA)-supplemented media.

**Figure 4:**
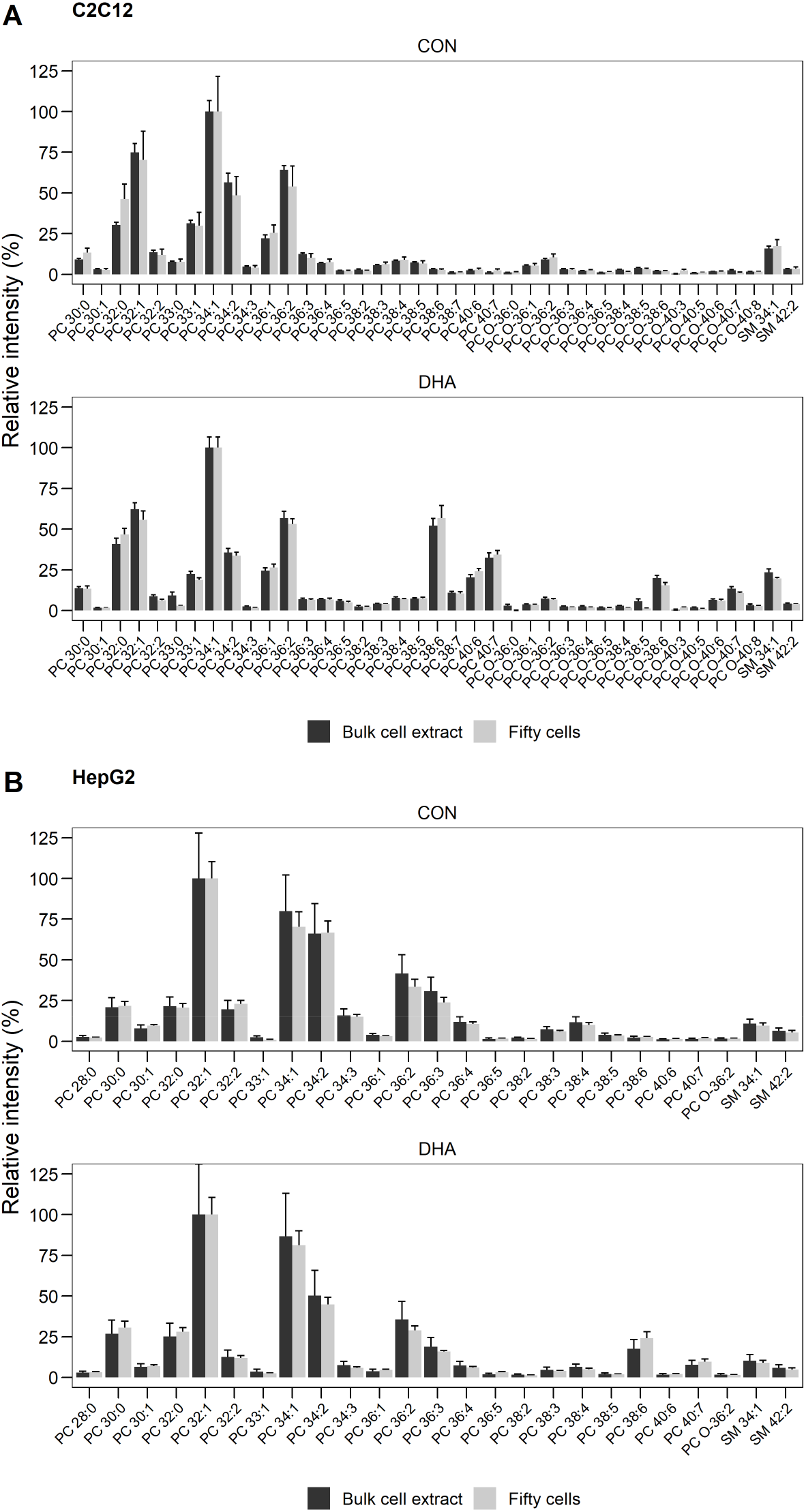
Comparison of phosphatidylcholine (PC) and sphingomyelin (SM) species detected in either bulk cell extract or fifty cells obtained by fluorescence-assisted cell sorting (FACS) for both (A) C2C12 cells and (B) HepG2 cells treated grown in control (CON) or docosahexaenoic acid (DHA)-supplemented media. Lipids shown are present at >1% relative abundance in either sample, values are mean±SEM (n=3). Welch t-test, * *P* < 0.05. Statistical output from all PC and SM species detected is available in the Supporting information (Table S2).

**Figure 5:**
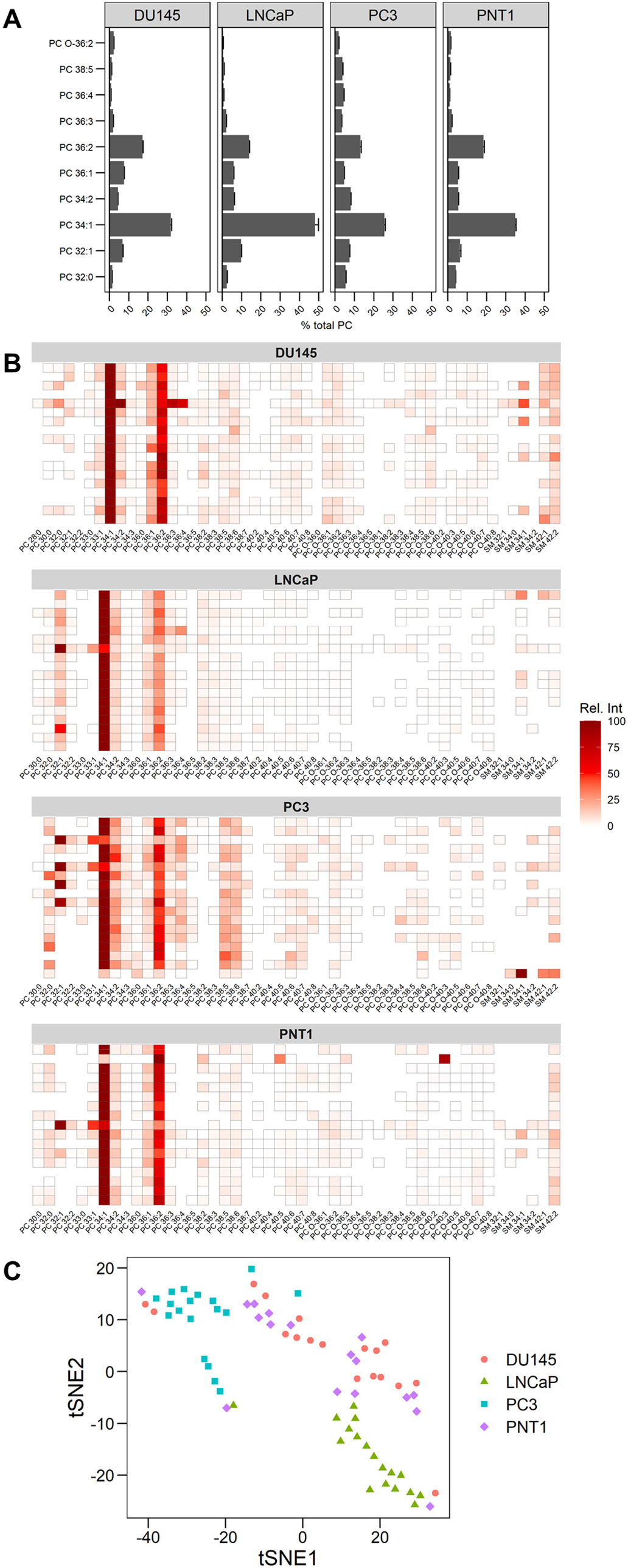
Analysis of phosphatidylcholine (PC) and sphingomyelin (SM) species from four different prostate cell lines, including cancerous (DU145, PC3 & LNCaP) and non-tumorigenic (PNT1) cells: (A) the ten most abundant PC species detected from the four prostate cell lines. Data was acquired from fifty cells obtained by fluorescence-assisted cell sorting (n=3) and expressed as a percent of total PC (±SEM). (B) PC and SM species detect ed from single cells across the four prostate cell lines (n=18). Data are expressed as relative intensity within each cell. (C) t-distributed stochastic neighbor embedding (tSNE) plot of the single-cell data acquired from the four prostate cell lines.

## Discussion

In the present study we have described a high-throughput single-cell lipidomics workflow that can detect and perform relative quantification on a high number of PC and SM species from single cells. To do this we have developed a workflow that combines single-cell FACS with automated chip-based nanoESI and shotgun lipidomics. This workflow utilizes a 96-well plate platform for both FACS and mass spectrometry but could be easily adapted to a 384-well plate format to further increase the throughput. We have used this workflow to study single-cell heterogeneity across two different cell lineages and have validated the lipids measured as being true biological measurements by performing experiments designed to manipulate the cell lipidome in a very predictable way (*i*.*e*., overnight DHA supplementation, Figures 1-3). Using this approach we could detect and quantify more than 50 different PC and SM species within singly-isolated C2C12 and HepG2 cells. We have also applied our method to the study of single-cell lipidome heterogeneity across four different prostate cell lines, including three cancerous (DU145, LNCaP, PC3) and one non-tumorigenic cell line (PNT1). Together these validate the robustness of the method and highlight its potential for detecting clinically relevant parameters.

FACS is a common method used for single-cell isolation and is routinely complexed with other single-cell profiling methodologies including transcriptomics. It has been previously reported that FACS can affect the polar metabolome of cells significantly (36); however, we have determined that it has little to no discernible impact on the PC and SM profile of cells (**Error! Reference source not found**.). Other single-cell isolation methods such as microfluidics, magnetic-activated cell sorting, and manual cell picking might also be easily complexed upstream of our single-cell shotgun lipidomics platform in place of FACS if desired. However, the use of FACS for single sorting offers the ability to sort cells based on cell surface markers to deliver enhanced information relevant to cell phenotypes. Modern FACS instruments can sort cells by up to 15 or more fluorescently labelled species per cell, which would allow the linking of complex cell phenotypes to their lipidome to provide enhanced detail on phenotype-driven heterogeneity (41). Moreover, when used in parallel with existing single-cell transcriptomics methods it could provide near-complete multi-omics data acquisition of distinct cell populations sharing phenotypic traits. In turn such data would provide a much more thorough understanding of the role of cellular heterogeneity across a range of fundamental biological and disease-driven research questions and could augment the delivery of personalized medicines for diseases with high levels of inherent lipid heterogeneity (*e*.*g*., certain cancers).

In this work we chose to apply our method to the study of prostate cancer for two main reasons. Firstly we know that prostate cancer cell lines differ in the abundance of discrete PC species present (37–39). Furthermore, changes in lipid metabolism have clinical significance in prostate cancer and are highly implicated in its pathogenesis, metastatic potential and resistance to treatment (39, 40). Profiling of PC and SM from singly isolated prostate cancer cell lines revealed both intra- and extra-cell line heterogeneity, and three broad groups of cells were able to be clustered by t-SNE: PC3, LNCaP and a third group of combined DU145 and PNT1 cells (**Error! Reference source not found**.C). This is an interesting finding as PNT1 cells are a non-tumorigenic immortalized prostate epithelial line (42), yet show a similar lipidome to CNS metastases-derived DU145 primary prostate adenocarcinoma cells (43). In terms of metastatic potential, PC3 cells are thought to be the most aggressive, with LNCaP being relatively more quiescent and DU145 somewhere in between (44, 45). The relative abundance of PC species detected within PC3, LNCaP and DU145 prostate cancer cells lines in the present study aligns well with that previously reported (37–39); however, we could find no publicly available datasets describing the lipidome of PNT1 prostate epithelial cells. Young et al. also described a similar PC profile between DU145 cells and the immortalized BPH-1 prostate epithelial cell line; however, significant differences were observed between phospholipid double-bond positional isomers and the expression of fatty acid desaturase 2 (FADS2) relative to stearoyl-CoA desaturase 1 (SCD1) (38). This translates to prostate cancer cell lines containing a different fatty acid carbon-carbon double bond profile when compared with prostate epithelial cells, but lipid carbon-carbon double bond position cannot be measured by traditional lipidomics profiling by tandem mass spectrometry. We also know that extracellular fatty acids heavily influence lipid metabolism in prostate cancer cells (46) and so the lipidome of our DU145 and PNT1 cells may simply represent the “default” or baseline lipid profile of prostate cells cultured *in vitro* in the absence of any influential biological effect. Future work may focus on complexing our single-cell lipidomics workflow with advanced mass spectrometry-based techniques that can determine more subtle lipid structural information. For example, ozone-induced dissociation could be used to determine the carbon-carbon double bond position in lipids to enhance the detail of data able to be collected from a single cell (47). The Paternò-Büchi reaction has also been used to determine carbon-carbon double bond position in single cells, but this reaction requires UV irradiation of the sample through a glass micropipette needle during electrospray and is likely not compatible with our workflow (25). Such data would likely be of increased clinical relevance in prostate and other cancers, and our method could easily be complexed with techniques such as liquid biopsy to study circulating tumor cells obtained from cancer patients. Excised tumors from patients could also feasibly be digested into single cells for measurement by our method, further adding to its potential applicability for cancer research.

While developing this workflow we also sought to expand coverage of the single-cell lipidome to include phosphatidylethanolamines (PE), which are typically the second-most abundant class of phospholipid present in mammalian cells. To do this we used a neutral ion loss scan of 141 Da in positive ion mode using settings previously reported (31). While we could detect numerous PE species in our bulk cell extracts and fifty FACS cell samples, significant contaminant species were detected within our single cell population including ions corresponding with PE 32:0 in both C2C12 and HepG2 cells and PE O-36:5 in HepG2 cells only (Supporting information, Figure S1). The overall relative intensity of ions detected using our neutral loss scan for PE was also significantly lower than that of PC/SM; leading us to conclude that PE was below the limit of detection for our single-cell method. Charge-switch derivatization strategies have been successfully used to improve detection in a range of lipid species (48, 49), and there is a chemical derivatization strategy that can impart a fixed positive charge on PE and phosphatidylserine aminophospholipids and increase sensitivity for such lipid species (50–53). Future work may target refining such derivatization strategies to expand lipidome coverage for single cells using our method. Moreover, our approach could be easily adapted to study the polar metabolome, and future work will focus on developing this methodology further. In particular, complexing our method to capillary electrophoresis may yield enhanced coverage and a high-throughput assay of the single-cell metabolome (54).

In summary, we have developed a high-throughput and quantitative method for the detection of PC and SM lipids from singly isolated cells. Using our method we could quantify more than 50 distinct PC and SM species from single cells of different cell lineages and observe inherent heterogeneity within similar cell populations but easily discriminate between different cell lines including the subtle variation in lipid profile present between prostate cancer cells. By using FACS to sort cells our method could be used to provide enhanced data on the relationship between cellular phenotypes and their lipidome, and our approach could be run in parallel with other single-cell methods to deliver near-complete multi-omics assay of cells of similar phenotypes. Our work continues to advance the progress of the development of single-cell lipidomics methods and has the capacity to improve our knowledge and understanding of cellular heterogeneity.

## Data availability

Data are able to be shared upon request to Sarah E Hancock

## Acknowledgements

We thank Dr Andrew Hoy from the University of Sydney for generously gifting us the prostate cell lines used in this work. The authors acknowledge use of the Mass Spectrometry Facility within School of Chemistry at the University of Wollongong.

## Abbreviations

(BHT): Butylated hydroxytoluene
(DHSM): dihydrosphingomyelin
(FCS): fetal calf serum
(FACS): fluorescence-assisted cell sorting
(LESA): liquid-extraction surface analysis
(nanoESI): nanoelectrospray ionization
(PBS): phosphate-buffered saline
(PC): phosphatidylcholine
(PE): phosphatidylethanolamine
(PUFA): polyunsaturated fatty acids
(S/N): signal-to-noise ratio
(SM): sphingomyelin
(t-SNE): t-distributed stochastic neighbor embedding

## References

1. Urban, P. L., T. Schmid, A. Amantonico, and R. Zenobi. 2011. Multidimensional Analysis of Single Algal Cells by Integrating Microspectroscopy with Mass Spectrometry. Anal. Chem. 83: 1843–1849.

2. Kompauer, M., S. Heiles, and B. Spengler. 2017. Atmospheric pressure MALDI mass spectrometry imaging of tissues and cells at 1.4-μm lateral resolution. Nat. Methods. 14: 90–96.

3. Dueñas, M. E., J. J. Essner, and Y. J. Lee. 2017. 3D MALDI Mass Spectrometry Imaging of a Single Cell: Spatial Mapping of Lipids in the Embryonic Development of Zebrafish. Sci. Rep. 7: 14946.

4. Yang, B., N. H. Patterson, T. Tsui, R. M. Caprioli, and J. L. Norris. 2018. Single-Cell Mass Spectrometry Reveals Changes in Lipid and Metabolite Expression in RAW 264.7 Cells upon Lipopolysaccharide Stimulation. J. Am. Soc. Mass Spectrom. 29: 1012–1020.

5. Xie, W., D. Gao, F. Jin, Y. Jiang, and H. Liu. 2015. Study of Phospholipids in Single Cells Using an Integrated Microfluidic Device Combined with Matrix-Assisted Laser Desorption/Ionization Mass Spectrometry. Anal. Chem. 87: 7052–7059.

6. Krismer, J., J. Sobek, R. F. Steinhoff, S. R. Fagerer, M. Pabst, and R. Zenobi. 2015. Screening of Chlamydomonas reinhardtii Populations with Single-Cell Resolution by Using a High-Throughput Microscale Sample Preparation for Matrix-Assisted Laser Desorption Ionization Mass Spectrometry. Appl. Environ. Microbiol. 81: 5546–5551.

7. Krismer, J., M. Tamminen, S. Fontana, R. Zenobi, and A. Narwani. 2017. Single-cell mass spectrometry reveals the importance of genetic diversity and plasticity for phenotypic variation in nitrogen-limited Chlamydomonas. ISME J. 11: 988–998.

8. Niehaus, M., J. Soltwisch, M. E. Belov, and K. Dreisewerd. 2019. Transmission-mode MALDI-2 mass spectrometry imaging of cells and tissues at subcellular resolution. Nat. Methods. 16: 925–931.

9. Bowman, A. P., J. F. J. Bogie, J. J. A. Hendriks, M. Haidar, M. Belov, R. M. A. Heeren, and S. R. Ellis. 2019. Evaluation of lipid coverage and high spatial resolution MALDI-imaging capabilities of oversampling combined with laser post-ionisation. Anal. Bioanal. Chem. [online] https://doi.org/10.1007/s00216-019-02290-3 (Accessed February 23, 2020).

10. Rappez, L., M. Stadler, S. Triana, R. M. Gathungu, K. Ovchinnikova, P. Phapale, M. Heikenwalder, and T. Alexandrov. 2021. SpaceM reveals metabolic states of single cells. Nat. Methods. 18: 799–805.

11. Szakal, C., K. Narayan, J. Fu, J. Lefman, and S. Subramaniam. 2011. Compositional Mapping of the Surface and Interior of Mammalian Cells at Submicrometer Resolution. Anal. Chem. 83: 1207–1213.

12. Fletcher, J. S., S. Rabbani, A. Henderson, P. Blenkinsopp, S. P. Thompson, N. P. Lockyer, and J. C. Vickerman. 2008. A New Dynamic in Mass Spectral Imaging of Single Biological Cells. Anal. Chem. 80: 9058–9064.

13. Passarelli, M. K., A. G. Ewing, and N. Winograd. 2013. Single-Cell Lipidomics: Characterizing and Imaging Lipids on the Surface of Individual Aplysia californica Neurons with Cluster Secondary Ion Mass Spectrometry. Anal. Chem. 85: 2231–2238.

14. Shrestha, B., P. Sripadi, B. R. Reschke, H. D. Henderson, M. J. Powell, S. A. Moody, and A. Vertes. 2014. Subcellular Metabolite and Lipid Analysis of Xenopus laevis Eggs by LAESI Mass Spectrometry. PLOS ONE. 9: e115173.

15. Shrestha, B., and A. Vertes. 2009. In Situ Metabolic Profiling of Single Cells by Laser Ablation Electrospray Ionization Mass Spectrometry. Anal. Chem. 81: 8265–8271.

16. Bergman, H.-M., and I. Lanekoff. 2017. Profiling and quantifying endogenous molecules in single cells using nano-DESI MS. Analyst. 142: 3639–3647.

17. Ellis, S. R., C. J. Ferris, K. J. Gilmore, T. W. Mitchell, S. J. Blanksby M., and in het Panhuis. 2012. Direct Lipid Profiling of Single Cells from Inkjet Printed Microarrays. Anal. Chem. 84: 9679–9683.

18. Snowden, S. G., H. J. R. Fernandes, J. Kent, S. Foskolou, P. Tate, S. F. Field, E. Metzakopian, and Koulman. 2020. Development and Application of High-Throughput Single Cell Lipid Profiling: A Study of SNCA-A53T Human Dopamine Neurons. iScience. 23. [online] https://www.cell.com/iscience/abstract/S2589-0042(20)30895-6 (Accessed November 12, 2020).

19. Phelps, M. S., and G. F. Verbeck. 2020. In Single Cell Metabolism: Methods and Protocols Methods in Molecular Biology (Shrestha, B., ed.). pp. 19–30., Springer, New York, NY. [online] https://doi.org/10.1007/978-1-4939-9831-9_3 (Accessed September 2, 2022).

20. Phelps, M. S., and G. F. Verbeck. 2015. A lipidomics demonstration of the importance of single cell analysis. Anal. Methods. 7: 3668–3670.

21. Pan, N., W. Rao, N. R. Kothapalli, R. Liu, A. W. G. Burgett, and Z. Yang. 2014. The Single-Probe: A Miniaturized Multifunctional Device for Single Cell Mass Spectrometry Analysis. Anal. Chem. 86: 9376–9380.

22. Sun, M., Z. Yang, and B. Wawrik. 2018. Metabolomic Fingerprints of Individual Algal Cells Using the Single-Probe Mass Spectrometry Technique. Front. Plant Sci. 9. [online] https://www.frontiersin.org/articles/10.3389/fpls.2018.00571/full (Accessed September 12, 2020).

23. Standke, S. J., D. H. Colby, R. C. Bensen, A. W. G. Burgett, and Z. Yang. 2019. Mass Spectrometry Measurement of Single Suspended Cells Using a Combined Cell Manipulation System and a Single-Probe Device. Anal. Chem. 91: 1738–1742.

24. Wang, R., H. Zhao, X. Zhang, X. Zhao, Z. Song, and J. Ouyang. 2019. Metabolic Discrimination of Breast Cancer Subtypes at the Single-Cell Level by Multiple Microextraction Coupled with Mass Spectrometry. Anal. Chem. 91: 3667–3674.

25. Zhu, Y., W. Wang, and Z. Yang. 2020. Combining Mass Spectrometry with Paternò–Büchi Reaction to Determine Double-Bond Positions in Lipids at the Single-Cell Level. Anal. Chem. [online] https://pubs.acs.org/doi/abs/10.1021/acs.analchem.0c02245 (Accessed February 10, 2022).

26. Sun, M., X. Chen, and Z. Yang. 2022. Single cell mass spectrometry studies reveal metabolomic features and potential mechanisms of drug-resistant cancer cell lines. Anal. Chim. Acta. 1206: 339761.

27. Chen, X., M. Sun, and Z. Yang. 2022. Single cell mass spectrometry analysis of drug-resistant cancer cells: Metabolomics studies of synergetic effect of combinational treatment. Anal. Chim. Acta. 1201: 339621.

28. Zhu, Y., R. Liu, and Z. Yang. 2019. Redesigning the T-probe for mass spectrometry analysis of online lysis of non-adherent single cells. Anal. Chim. Acta. 1084: 53–59.

29. Ellis, S. R., J. R. Hughes, T. W. Mitchell, M. in het Panhuis, and S. J. Blanksby. 2012. Using ambient ozone for assignment of double bond position in unsaturated lipids. Analyst. 137: 1100–1110.

30. Alsabeeh, N., B. Chausse, P. A. Kakimoto, A. J. Kowaltowski, and O. Shirihai. 2018. Cell Culture Models of Fatty Acid Overload: Problems and Solutions. Biochim. Biophys. Acta. 1863: 143–151.

31. Norris, S. E., M. G. Friedrich, T. W. Mitchell, R. J. W. Truscott, and P. L. Else. 2015. Human prefrontal cortex phospholipids containing docosahexaenoic acid increase during normal adult aging, whereas those containing arachidonic acid decrease. Neurobiol. Aging. 36: 1659–69.

32. R Core Team. R: A language and environment for statistical computing.

33. Liebisch, G., J. A. Vizcaíno, H. Köfeler, M. Trötzmüller, W. J. Griffiths, G. Schmitz, F. Spener, and M. J. O. Wakelam. 2013. Shorthand notation for lipid structures derived from mass spectrometry. J. Lipid Res. 54: 1523–1530.

34. Krijthe, J. J. 2015. Rtsne: T-Distributed Stochastic Neighbor Embedding using a Barnes-Hut Implementation. [online] https://github.com/jkrijthe/Rtsne.

35. Burdge, G. C., and P. C. Calder. 2005. Conversion of α-linolenic acid to longer-chain polyunsaturated fatty acids in human adults. Reprod. Nutr. Dev. 45: 581–597.

36. Binek, A., D. Rojo, J. Godzien, F. J. Rupérez, V. Nuñez, I. Jorge, M. Ricote, J. Vázquez, and C. Barbas. 2019. Flow Cytometry Has a Significant Impact on the Cellular Metabolome. J. Proteome Res. 18: 169–181.

37. Sorvina, A., C. A. Bader, C. Caporale, E. A. Carter, I. R. D. Johnson, E. J. Parkinson-Lawrence, P. V. Simpson, P. J. Wright, S. Stagni, P. A. Lay, M. Massi, D. A. Brooks, and S. E. Plush. 2018. Lipid profiles of prostate cancer cells. Oncotarget. 9: 35541–35552.

38. Young, R. S. E., A. P. Bowman, E. D. Williams, K. D. Tousignant, C. L. Bidgood, V. R. Narreddula, R. Gupta, D. L. Marshall, B. L. J. Poad, C. C. Nelson, S. R. Ellis, R. M. A. Heeren, M. C. Sadowski, and S. J. Blanksby. 2021. Apocryphal FADS2 activity promotes fatty acid diversification in cancer. Cell Rep. 34. [online] https://www.cell.com/cell-reports/abstract/S2211-1247(21)00051-6 (Accessed February 16, 2021).

39. Ingram, L. M., M. C. Finnerty, M. Mansoura, C.-W. Chou, and B. S. Cummings. 2021. Identification of lipidomic profiles associated with drug-resistant prostate cancer cells. Lipids Health Dis. 20: 15.

40. Butler, L. M., Y. Perone, J. Dehairs, L. E. Lupien, V. de Laat, A. Talebi, M. Loda, W. B. Kinlaw, and J. V. Swinnen. 2020. Lipids and cancer: Emerging roles in pathogenesis, diagnosis and therapeutic intervention. Adv. Drug Deliv. Rev. 159: 245–293.

41. McKinnon, K. M. 2018. Flow Cytometry: An Overview. Curr. Protoc. Immunol. 120: 5.1.1-5.1.11.

42. Cussenot, O., P. Berthon, R. Berger, I. Mowszowicz, A. Faille, F. Hojman, P. Teillac, A. Le Duc, and F. Calvo. 1991. Immortalization of human adult normal prostatic epithelial cells by liposomes containing large T-SV40 gene. J. Urol. 146: 881–886.

43. Stone, K. R., D. D. Mickey, H. Wunderli, G. H. Mickey, and D. F. Paulson. 1978. Isolation of a human prostate carcinoma cell line (DU 145). Int. J. Cancer. 21: 274–281.

44. Tai, S., Y. Sun, J. M. Squires, H. Zhang, W. K. Oh, C.-Z. Liang, and J. Huang. 2011. PC3 Is a Cell Line Characteristic of Prostatic Small Cell Carcinoma. The Prostate. 71: 1668–1679.

45. Webber, M. M., D. Bello, and S. Quader. 1997. Immortalized and tumorigenic adult human prostatic epithelial cell lines: Characteristics and applications part 2. Tumorigenic cell lines. The Prostate. 30: 58–64.

46. Balaban, S., Z. D. Nassar, A. Y. Zhang, E. Hosseini-Beheshti, M. M. Centenera, M. Schreuder, H.-M. Lin, A. Aishah, B. Varney, F. Liu-Fu, L. S. Lee, S. R. Nagarajan, R. F. Shearer, R.-A. Hardie, N. L. Raftopulos, M. S. Kakani, D. N. Saunders, J. Holst, L. G. Horvath, L. M. Butler, and A. J. Hoy. 2019. Extracellular Fatty Acids Are the Major Contributor to Lipid Synthesis in Prostate Cancer. Mol. Cancer Res. 17: 949–962.

47. Brown, S. H. J., T. W. Mitchell, and S. J. Blanksby. 2011. Analysis of unsaturated lipids by ozone-induced dissociation. Biochim. Biophys. Acta BBA - Mol. Cell Biol. Lipids. 1811: 807–817.

48. Bollinger, J. G., W. Thompson, Y. Lai, R. C. Oslund, T. S. Hallstrand, M. Sadilek, F. Turecek, and M. H. Gelb. 2010. Improved Sensitivity Mass Spectrometric Detection of Eicosanoids by Charge Reversal Derivatization. Anal. Chem. 82: 6790–6796.

49. Hancock, S. E., R. Ailuri, D. L. Marshall, S. H. J. Brown, J. T. Saville, V. R. Narreddula, N. R. Boase, B. L. J. Poad, A. J. Trevitt, M. D. P. Willcox, M. J. Kelso, T. W. Mitchell, and S. J. Blanksby. 2018. Mass spectrometry-directed structure elucidation and total synthesis of ultra-long chain (O-acyl)-ω-hydroxy fatty acids. J. Lipid Res. 59: 1510–1518.

50. Fhaner, C. J., S. Liu, X. Zhou, and G. E. Reid. 2013. Functional Group Selective Derivatization and Gas-Phase Fragmentation Reactions of Plasmalogen Glycerophospholipids. Mass Spectrom. 2. [online] http://www.ncbi.nlm.nih.gov/pmc/articles/PMC3810100/ (Accessed February 3, 2015).

51. Fhaner, C. J., S. Liu, H. Ji, R. J. Simpson, and G. E. Reid. 2012. Comprehensive Lipidome Profiling of Isogenic Primary and Metastatic Colon Adenocarcinoma Cell Lines. Anal. Chem. 84: 8917–8926.

52. Nie, S., C. J. Fhaner, S. Liu, D. Peake, R. Kiyonami, Y. Huang, and G. E. Reid. 2015. Characterization and multiplexed quantification of derivatized aminophospholipids. Int. J. Mass Spectrom. 391: 71–81.

53. Ryan, E., and G. E. Reid. 2016. Chemical Derivatization and Ultrahigh Resolution and Accurate Mass Spectrometry Strategies for “Shotgun” Lipidome Analysis. Acc. Chem. Res. 49: 1596–1604.

54. Kawai, T., N. Ota, K. Okada, A. Imasato, Y. Owa, M. Morita, M. Tada, and Y. Tanaka. 2019. Ultrasensitive Single Cell Metabolomics by Capillary Electrophoresis–Mass Spectrometry with a Thin-Walled Tapered Emitter and Large-Volume Dual Sample Preconcentration. Anal. Chem. 91: 10564–10572.

