## Supporting information for "FACS-assisted single-cell lipidome analysis of phosphatidylcholines and sphingomyelins"

.

#### Tables

**Table S1:** Target list used for lipid identification in Lipidview software.

| Name | Mass (m/z) | Precursor /NL of (Da) | Ion mode | Scan type | Class | Species | IS name | Isotopic correction factor |
|---|---|---|---|---|---|---|---|---|
| Reference Peak Information | | | | | | | | |
| IS PC 17:0/17:0 | 762.5 | 184.1 | Positive | PC | | | PC 34:0 | 1.642 |
| IS DHSM 12:0 | 649.5 | 184.1 | Positive | SM/PC | | | DHSM 12:0 | 1.516 |
| IS PE 17:0/17:0 | 720.5 | 141 | Positive | -PE | | | PE 34:0 | 1.588 |
| Target Lipid Information | | | | | | | | |
| PC 28:0 | 678.6 | 184.1 | Positive | SM/PC | PC | PC 28:0 | PC 34:0 | 1.535 |
| PC 30:1 | 704.6 | 184.1 | Positive | SM/PC | PC | PC 30:1 | PC 34:0 | 1.569 |
| PC 30:0 | 706.6 | 184.1 | Positive | SM/PC | PC | PC 30:0 | PC 34:0 | 1.57 |
| PC 32:2 | 730.6 | 184.1 | Positive | SM/PC | PC | PC 32:2 | PC 34:0 | 1.605 |
| PC 32:1 | 732.6 | 184.1 | Positive | SM/PC | PC | PC 32:1 | PC 34:0 | 1.605 |
| PC 32:0 | 734.6 | 184.1 | Positive | SM/PC | PC | PC 32:0 | PC 34:0 | 1.605 |
| PC 34:4 | 754.6 | 184.1 | Positive | SM/PC | PC | PC 34:4 | PC 34:0 | 1.641 |
| PC 34:3 | 756.6 | 184.1 | Positive | SM/PC | PC | PC 34:3 | PC 34:0 | 1.641 |
| PC 34:2 | 758.7 | 184.1 | Positive | SM/PC | PC | PC 34:2 | PC 34:0 | 1.641 |
| PC 34:1 | 760.6 | 184.1 | Positive | SM/PC | PC | PC 34:1 | PC 34:0 | 1.642 |
| PC 33:1 | 746.6 | 184.1 | Positive | SM/PC | PC | PC 33:1 | PC 34:0 | 1.64 |
| PC 33:0 | 744.7 | 184.1 | Positive | SM/PC | PC | PC 33:0 | PC 34:0 | 1.64 |
| PC O-36:5 | 766.7 | 184.1 | Positive | SM/PC | PC | PC O-36:5 | PC 34:0 | 1.674 |
| PC O-36:4 | 768.7 | 184.1 | Positive | SM/PC | PC | PC O-36:4 | PC 34:0 | 1.674 |
| PC O-36:3 | 770.6 | 184.1 | Positive | SM/PC | PC | PC O-36:3 | PC 34:0 | 1.675 |
| PC O-36:2 | 772.3 | 184.1 | Positive | SM/PC | PC | PC O-36:2 | PC 34:0 | 1.675 |
| PC O-36:1 | 774.4 | 184.1 | Positive | SM/PC | PC | PC O-36:1 | PC 34:0 | 1.675 |
| PC O-36:0 | 776.6 | 184.1 | Positive | SM/PC | PC | PC O-36:0 | PC 34:0 | 1.676 |
| PC 36:5 | 780.6 | 184.1 | Positive | SM/PC | PC | PC 36:5 | PC 34:0 | 1.678 |
| PC 36:4 | 782.6 | 184.1 | Positive | SM/PC | PC | PC 36:4 | PC 34:0 | 1.678 |
| PC 36:3 | 784.6 | 184.1 | Positive | SM/PC | PC | PC 36:3 | PC 34:0 | 1.678 |
| PC 36:2 | 786.6 | 184.1 | Positive | SM/PC | PC | PC 36:2 | PC 34:0 | 1.679 |
| PC 36:1 | 788.7 | 184.1 | Positive | SM/PC | PC | PC 36:1 | PC 34:0 | 1.679 |
| PC 36:0 | 790.3 | 184.1 | Positive | SM/PC | PC | PC 36:0 | PC 34:0 | 1.679 |
| PC O-38:6 | 792.4 | 184.1 | Positive | SM/PC | PC | PC O-38:6 | PC 34:0 | 1.712 |
| PC O-38:5 | 794.4 | 184.1 | Positive | SM/PC | PC | PC O-38:5 | PC 34:0 | 1.712 |
| PC O-38:4 | 796.4 | 184.1 | Positive | SM/PC | PC | PC O-38:4 | PC 34:0 | 1.712 |
| PC O-38:3 | 798.5 | 184.1 | Positive | SM/PC | PC | PC O-38:3 | PC 34:0 | 1.713 |
| PC O-38:2 | 800.8 | 184.1 | Positive | SM/PC | PC | PC O-38:2 | PC 34:0 | 1.713 |
| PC O-38:1 | 802.8 | 184.1 | Positive | SM/PC | PC | PC 38:8 | PC 34:0 | 1.715 |
| PC 38:7 | 804.6 | 184.1 | Positive | SM/PC | PC | PC 38:7 | PC 34:0 | 1.715 |
| PC 38:6 | 806.6 | 184.1 | Positive | SM/PC | PC | PC 38:6 | PC 34:0 | 1.715 |
| PC 38:5 | 808.6 | 184.1 | Positive | SM/PC | PC | PC 38:5 | PC 34:0 | 1.716 |
| PC 38:4 | 810.6 | 184.1 | Positive | SM/PC | PC | PC 38:4 | PC 34:0 | 1.716 |
| PC 38:3 | 812.7 | 184.1 | Positive | SM/PC | PC | PC 38:3 | PC 34:0 | 1.716 |

| | | | | | | | | |
|---|---|---|---|---|---|---|---|---|
| PC 38:2 | 814.7 | 184.1 | Positive | SM/PC | PC | PC 38:2 | PC 34:0 | 1.717 |
| PC O-40:8 | 816.7 | 184.1 | Positive | SM/PC | PC | PC O-40:8 | PC 34:0 | 1.75 |
| PC O-40:7 | 818.7 | 184.1 | Positive | SM/PC | PC | PC O-40:7 | PC 34:0 | 1.75 |
| PC O-40:6 | 820.7 | 184.1 | Positive | SM/PC | PC | PC O-40:6 | PC 34:0 | 1.751 |
| PC O-40:5 | 822.7 | 184.1 | Positive | SM/PC | PC | PC O-40:5 | PC 34:0 | 1.751 |
| PC O-40:4 | 824.7 | 184.1 | Positive | SM/PC | PC | PC O-40:4 | PC 34:0 | 1.751 |
| PC O-40:3 | 826.4 | 184.1 | Positive | SM/PC | PC | PC O-40:3 | PC 34:0 | 1.752 |
| PC O-40:2 | 828.7 | 184.1 | Positive | SM/PC | PC | PC O-40:2 | PC 34:0 | 1.752 |
| PC 40:8 | 830.7 | 184.1 | Positive | SM/PC | PC | PC 40:8 | PC 34:0 | 1.752 |
| PC 40:7 | 832.5 | 184.1 | Positive | SM/PC | PC | PC 40:7 | PC 34:0 | 1.753 |
| PC 40:6 | 834.7 | 184.1 | Positive | SM/PC | PC | PC 40:6 | PC 34:0 | 1.755 |
| PC 40:5 | 836.7 | 184.1 | Positive | SM/PC | PC | PC 40:5 | PC 34:0 | 1.755 |
| PC 40:4 | 838.5 | 184.1 | Positive | SM/PC | PC | PC 40:4 | PC 34:0 | 1.755 |
| PC 40:3 | 840.6 | 184.1 | Positive | SM/PC | PC | PC 40:3 | PC 34:0 | 1.756 |
| PC 40:2 | 842.5 | 184.1 | Positive | SM/PC | PC | PC 40:2 | PC 34:0 | 1.756 |
| SM 32:1;2 | 675.6 | 184.1 | Positive | SM/PC | SM | SM 32:1;2 | DHSM 12:0 | 1.55 |
| SM 34:2;2 | 701.6 | 184.1 | Positive | SM/PC | SM | SM 34:2;2 | DHSM 12:0 | 1.586 |
| SM 34:1;2 | 703.6 | 184.1 | Positive | SM/PC | SM | SM 34:1;2 | DHSM 12:0 | 1.586 |
| SM 34:0;2 | 705.6 | 184.1 | Positive | SM/PC | SM | SM 34:0;2 | DHSM 12:0 | 1.586 |
| SM 42:2;2 | 813.6 | 184.1 | Positive | SM/PC | SM | SM 42:2;2 | DHSM 12:0 | 1.736 |
| SM 42:1;2 | 815.6 | 184.1 | Positive | SM/PC | SM | SM 42:1;2 | DHSM 12:0 | 1.736 |
| PE 32:2 | 688.7 | 141 | Positive | -PE | PE | PE 32:2 | PE 34:0 | 1.551 |
| PE 32:1 | 690.7 | 141 | Positive | -PE | PE | PE 32:1 | PE 34:0 | 1.552 |
| PE 32:0 | 692.7 | 141 | Positive | -PE | PE | PE 32:0 | PE 34:0 | 1.552 |
| PE 34:3 | 714.6 | 141 | Positive | -PE | PE | PE 34:3 | PE 34:0 | 1.586 |
| PE 34:2 | 716.6 | 141 | Positive | -PE | PE | PE 34:2 | PE 34:0 | 1.587 |
| PE 34:1 | 718.6 | 141 | Positive | -PE | PE | PE 34:1 | PE 34:0 | 1.587 |
| PE O-36:5 | 724.6 | 141 | Positive | -PE | PE | PE O-36:5 | PE 34:0 | 1.6183 |
| PE 36:6 | 736.6 | 141 | Positive | -PE | PE | PE 36:6 | PE 34:0 | 1.622 |
| PE 36:5 | 738.6 | 141 | Positive | -PE | PE | PE 36:5 | PE 34:0 | 1.622 |
| PE 36:4 | 740.7 | 141 | Positive | -PE | PE | PE 36:4 | PE 34:0 | 1.622 |
| PE 36:3 | 742.6 | 141 | Positive | -PE | PE | PE 36:3 | PE 34:0 | 1.623 |
| PE 36:2 | 744.6 | 141 | Positive | -PE | PE | PE 36:2 | PE 34:0 | 1.623 |
| PE 36:1 | 746.7 | 141 | Positive | -PE | PE | PE 36:1 | PE 34:0 | 1.623 |
| PE 36:0 | 748.6 | 141 | Positive | -PE | PE | PE 36:0 | PE 34:0 | 1.624 |
| PE O-38:6 | 750.6 | 141 | Positive | -PE | PE | PE O-38:6 | PE 34:0 | 1.6549 |
| PE O-38:2 | 758.8 | 141 | Positive | -PE | PE | PE O-38:2 | PE 34:0 | 1.6562 |
| PE 38:7 | 762.5 | 141 | Positive | -PE | PE | PE 38:7 | PE 34:0 | 1.658 |
| PE 38:6 | 764.6 | 141 | Positive | -PE | PE | PE 38:6 | PE 34:0 | 1.658 |
| PE 38:5 | 766.6 | 141 | Positive | -PE | PE | PE 38:5 | PE 34:0 | 1.659 |
| PE 38:4 | 768.6 | 141 | Positive | -PE | PE | PE 38:4 | PE 34:0 | 1.659 |
| PE 38:3 | 770.6 | 141 | Positive | -PE | PE | PE 38:3 | PE 34:0 | 1.659 |
| PE 38:2 | 772.7 | 141 | Positive | -PE | PE | PE 38:2 | PE 34:0 | 1.66 |
| PE O-40:2 | 786.6 | 141 | Positive | -PE | PE | PE O-40:2 | PE 34:0 | 1.6939 |
| PE 40:8 | 788.6 | 141 | Positive | -PE | PE | PE 40:8 | PE 34:0 | 1.696 |
| PE 40:7 | 790.6 | 141 | Positive | -PE | PE | PE 40:7 | PE 34:0 | 1.696 |

| | | | | | | | | |
|---|---|---|---|---|---|---|---|---|
| PE 40:6 | 792.6 | 141 | Positive | -PE | PE | PE 40:6 | PE 34:0 | 1.696 |
| PE 40:5 | 794.6 | 141 | Positive | -PE | PE | PE 40:5 | PE 34:0 | 1.697 |
| PE 40:2 | 800.8 | 141 | Positive | -PE | PE | PE 40:2 | PE 34:0 | 1.698 |
| PE O-42:4 | 810.5 | 141 | Positive | -PE | PE | PE 41:4 | PE 34:0 | 1.7318 |
| PE O-42:2 | 814.6 | 141 | Positive | -PE | PE | PE 41:2 | PE 34:0 | 1.7325 |

*DHSM* dihydrosphingomyelin, *IS* internal standard, *NL* neutral loss, *PC* phosphatidylcholine, *PE* phosphatidylethanolamine

**Table S2:** Statistical output from all detected PC and SM species significantly different between bulk cell extracts and fifty cells obtained by FACS for both C2C12 and HepG2 cells grown in control or DHA-supplemented media

| | | | Mean values* | | | | | | | |
|---|---|---|---|---|---|---|---|---|---|---|
| **Cell line** | **Lipid species** | **Media** | **Bulk cell extract** | **Fifty cells** | **Difference** | **statistic** | **p.value** | **parameter** | **conf.low** | **conf.high** |
| C2C12 | PC 28:0 | CON | 0.677267 | 0.057315 | 0.619952 | 7.711064 | 0.007596 | 2.576296 | 0.338557 | 0.901348 |
| C2C12 | PC 40:5 | DHA | 1.136988 | 0.470475 | 0.666513 | 4.506505 | 0.027431 | 2.604082 | 0.152637 | 1.180389 |
| C2C12 | PC O-38:3 | CON | 1.503391 | 0.319038 | 1.184353 | 4.514423 | 0.018057 | 3.16842 | 0.373996 | 1.99471 |
| C2C12 | PC O-38:4 | DHA | 2.905955 | 1.597656 | 1.308299 | 3.842199 | 0.029799 | 3.07324 | 0.239115 | 2.377482 |
| C2C12 | PC O-40:3 | DHA | 0.45247 | 1.892797 | -1.44033 | -3.80111 | 0.02208 | 3.692901 | -2.5278 | -0.35285 |
| C2C12 | PC O-40:5 | DHA | 1.854475 | 1.109687 | 0.744788 | 3.188368 | 0.042093 | 3.383427 | 0.046844 | 1.442733 |
| C2C12 | SM 34:0 | CON | 1.533367 | 0.051599 | 1.481768 | 5.949405 | 0.022071 | 2.17902 | 0.490243 | 2.473292 |
| C2C12 | SM 34:0 | DHA | 0.948997 | 0.218288 | 0.730709 | 3.42506 | 0.039521 | 3.106405 | 0.064729 | 1.39669 |
| C2C12 | SM 34:2 | CON | 1.819516 | 0 | 1.819516 | 22.48913 | 0.001971 | 2 | 1.471403 | 2.167628 |
| C2C12 | SM 34:2 | DHA | 2.868861 | 0.659312 | 2.20955 | 5.528401 | 0.026605 | 2.146535 | 0.597638 | 3.821462 |
| HepG2 | PC 33:0 | CON | 0 | 1.173872 | -1.17387 | -11.9612 | 0.006917 | 2 | -1.59613 | -0.75161 |
| HepG2 | PC O-36:0 | CON | 0 | 0.263927 | -0.26393 | -8.45971 | 0.013687 | 2 | -0.39816 | -0.12969 |
| HepG2 | PC O-36:5 | DHA | 0.037858 | 0.307606 | -0.26975 | -9.96044 | 6.67E-04 | 3.875572 | -0.3459 | -0.19359 |
| HepG2 | PC O-38:4 | CON | 0.908009 | 0.209331 | 0.698678 | 3.284398 | 0.042226 | 3.195523 | 0.044529 | 1.352827 |
| HepG2 | PC O-40:2 | CON | 0.011522 | 0.583806 | -0.57228 | -7.47863 | 0.015332 | 2.092745 | -0.88794 | -0.25663 |
| HepG2 | PC O-40:3 | CON | 0.181543 | 1.176038 | -0.9945 | -5.11354 | 0.012925 | 3.140889 | -1.59801 | -0.39098 |
| HepG2 | PC O-40:5 | CON | 0.005649 | 0.280797 | -0.27515 | -19.382 | 5.31E-04 | 2.726945 | -0.323 | -0.2273 |
| HepG2 | PC O-40:5 | DHA | 9.94E-04 | 0.256989 | -0.256 | -6.05471 | 0.026142 | 2.002211 | -0.43772 | -0.07427 |
| HepG2 | SM 32:1 | CON | 1.193725 | 0.286911 | 0.906814 | 3.49325 | 0.041589 | 2.910934 | 0.066196 | 1.747433 |
| HepG2 | SM 34:2 | CON | 0.652823 | 0 | 0.652823 | 11.34558 | 0.007679 | 2 | 0.405249 | 0.900397 |

*Mean values calculated from the relative abundance of PC & SM species (n=3). Welch t test, $P < 0.05$

### Figures

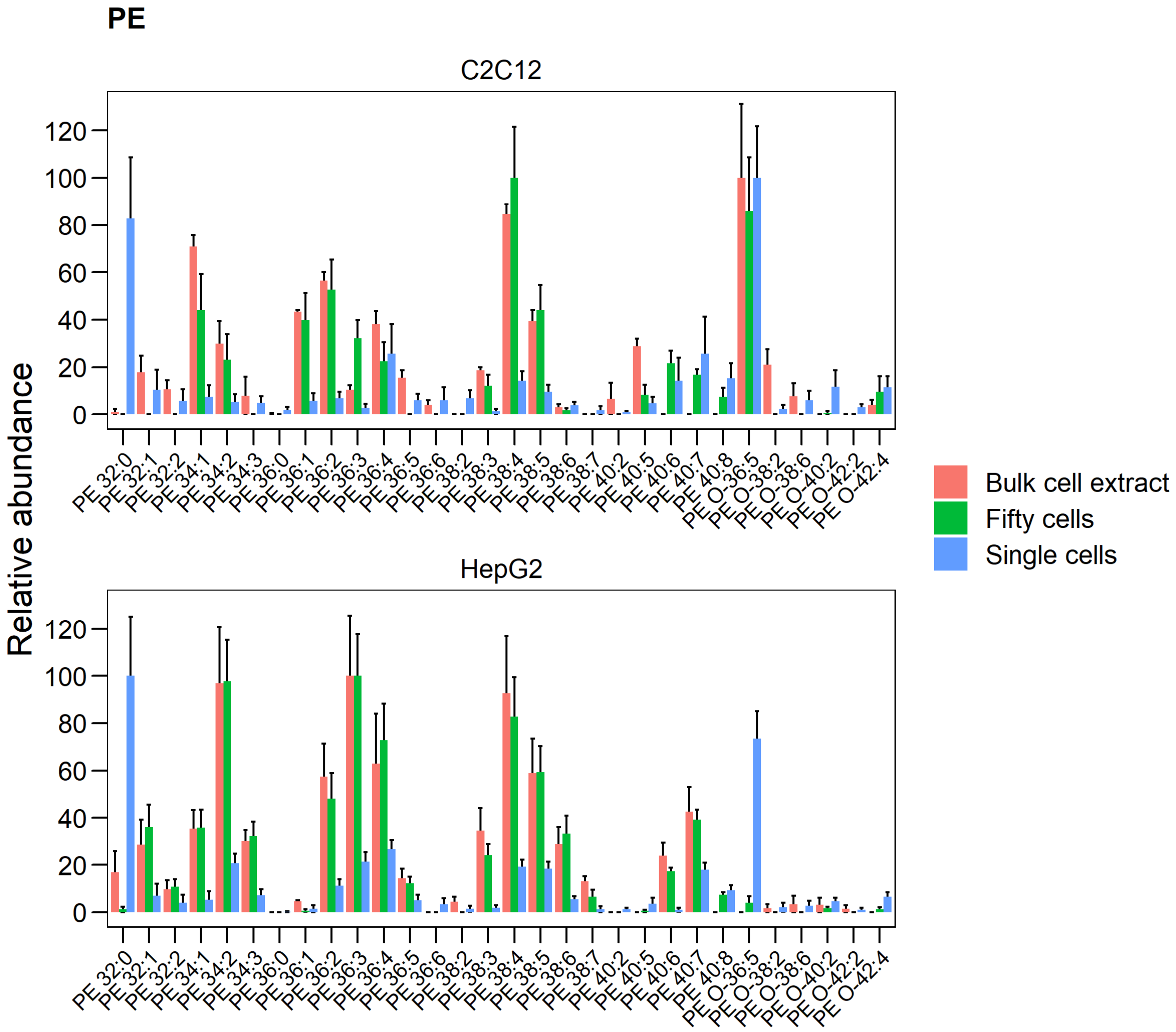


Figure S1: relative abundance of phosphatidylethanolamine (PE) species detected from bulk cell extracts (n=3), fifty sorted cells (n=3) and single sorted cells (n=17-18) by shotgun lipidomics. Values are mean(±SEM).
